# The antimicrobial metabolite nisin Z reduces intestinal tumorigenesis and modulates the cecal microbiome in Apc^Min/+^ mice

**DOI:** 10.1101/2025.05.18.654755

**Authors:** Layan Hamidi Nia, Sara Alqudah, Rachel L. Markley, Beckey DeLucia, Viharika Bobba, Judi Elmallah, Ina Nemet, Naseer Sangwan, Jan Claesen

## Abstract

Nisin Z, an antimicrobial metabolite produced by *Lactococcus lactis spp.,* has been safely used as a food preservative for many years. Nisin Z also showed promising activity against various cancer types *in vitro*, and significantly reduced tumor size in an ectopic head and neck cancer model. Here, we investigate the activity of nisin Z for colorectal cancer treatment and observed an *in vitro* reduction in cellular proliferation, and a moderate enhancement in cell death. We next analyzed the effect of oral nisin Z administration in the Apc^min/+^ intestinal adenoma mouse model. We measured tumor burden along the gastrointestinal tract and observed a decrease in tumor burden in the middle region of the small intestine, but not in the lower region or colon. Since tumor progression in the Apc^min/+^ model is exacerbated by an inflammatory environment, we next determined whether nisin Z impacts this in a direct or indirect manner. We show that nisin Z can directly reduce NF-κB activation in a dose-dependent manner. In addition, nisin Z impacted the cecal microbiome composition as well as microbiota-associated plasma metabolites, causing an overall shift towards a more health-associated profile. Interestingly, the Apc^min/+^ genotype differentially impacted the nisin Z-mediated differences in cecal microbiome composition and plasma metabolites compared to wildtype animals. In summary, our data suggest that the reduction in small intestinal tumor burden could be due to nisin Z’s contribution to a reduced pro-inflammatory environment. Future studies will reveal whether nisin’s localized effect is due to degradation of the peptidic compound in more distal regions of the gastrointestinal tract and focus on development of delivery systems to increase efficacy.

**Importance:** With the increased incidence of colorectal cancer, especially among younger individuals, it is critical to study approaches that help with the prevention and treatment of this debilitating disease. Our study indicates that nisin Z, a bacterially produced peptide antibiotic, decreases the growth of colorectal cancer cells and moderately increases cell death *in vitro*. Oral administration of nisin Z in an intestinal adenoma mouse model revealed a reduction of tumor burden in the middle region of the small intestine. This decreased tumor burden might in part be attributed to a direct anti-inflammatory effect, as well as an indirect effect on the gut microbiota and their metabolites due to nisin Z’s antibacterial activity. Overall, we demonstrate a potential activity for nisin Z in the prevention or amelioration of inflammation-associated colorectal cancer, underscoring the significance of investigating the properties of bacterial natural products in human health.

## Introduction

Colorectal cancer (CRC) is the third most common cancer type worldwide, typically affecting individuals over the age of 50. CRC can have genetic origins, causing an increased predilection for development of polyps in the colonic or rectal epithelium^1,2^. In addition, CRC progression can be influenced by a combination of factors, including lifestyle choices and genetic conditions^3,4^. The gut microbiota can play a crucial role in regulating CRC through various microbiome-dependent metabolites. For example, indoles resulting from gut microbial tryptophan metabolism play an important role in maintaining intestinal barrier homeostasis and attenuating CRC progression^5^.

A common genetic mutation in the APC gene can lead to familial adenomatous polyposis, causing people to sporadically develop colorectal adenomas^5^. These characteristics are modeled by the Apc^Min/+^ mouse model, which predisposes mice to develop multiple intestinal neoplasia (min), resulting in tumor formation across their small intestine and colon^6^. The Apc^Min/+^ model is readily amenable to testing medicinal interventions as well as lifestyle choices aimed at the prevention of CRC development. As an example, the consumption of a highly processed and high-fat diet can lead to increased CRC development, which is paired with a dysbiosis of the gut microbiome^4^.

Nisin is a post-translationally modified peptide produced by *Lactococcus lactis* that has antibacterial properties against Gram-positive bacteria. Due to its antibiotic activity, nisin Z is commonly used as a food preservative in dairy products and has received the “Generally Recognized as Safe” (GRAS) status by the U.S. Food and Drug Administration (FDA). More recently, nisin and its variants have gained attention because of their potential use in medical applications, including gastrointestinal (GI) infections and treatment of certain cancer types^7,8,9,10,11^. Most notably, nisin Z has potent activity in reducing head and neck cancer cell proliferation as well as angiogenesis^9^. This effect has in part been attributed to increased calcium influx and subsequent cell cycle arrest mediated by the pro-apoptotic cation transport regulator CHAC1^12^. CHAC1 upregulation has been associated with various cancer types, including CRC, Orally administered nisin Z significantly reduces tumor proliferation in a preclinical model^9^, and is currently in a phase I clinical trial for squamous and head and neck carcinoma^13^.

In this paper, we will determine the impact of nisin Z on cellular proliferation, inflammatory pathways, and the gut microbiome, as it affects CRC. We found that nisin Z decreases colorectal cancer cellular proliferation while slightly enhancing cell death. We also show a dose-dependent effect on NF-κB activation, as well as shifts in the gut microbiome and microbial metabolites due to nisin Z’s antibacterial properties. .

## Results

### Nisin Z reduces Caco-2 cell proliferation and enhances cell death

Given the promising effects of nisin Z in attenuating head and neck cancer^9^, we investigated its potential activity on colorectal cancer. We first looked for the effect of nisin Z on Caco-2 (human colon adenocarcinoma) cell proliferation. Using an automated microplate live-cell imaging system, we monitored cellular confluency for 72 h at nisin Z concentrations up to 300 µM (1 mg/ml) (Fig. 1A, B). At concentrations of 30 µM or 150 µM, nisin Z treatment did not significantly affect the increase in Caco-2 confluency compared to the vehicle control, which reached ∼80% confluency at 72 h. However, at a nisin Z concentration of 300 µM, we observed a significant reduction in the cellular confluency increase after 48 h. To investigate whether this reduced confluency increase is driven by a decreased proliferation rate or increased cell death, we next performed a LIVE/DEAD stain to test for potential cytotoxicity after 24 h and 72 h nisin Z treatment (Fig. 1C). We observed a significant increase in percentage of dead cells after exposure to 300 µM nisin Z at 72 h. In summary, these results indicate that nisin Z treatment dose-dependently affects Caco-2 cell growth, which could be the result of an antiproliferative activity combined with a modest cytotoxic activity.

**Fig. 1.**
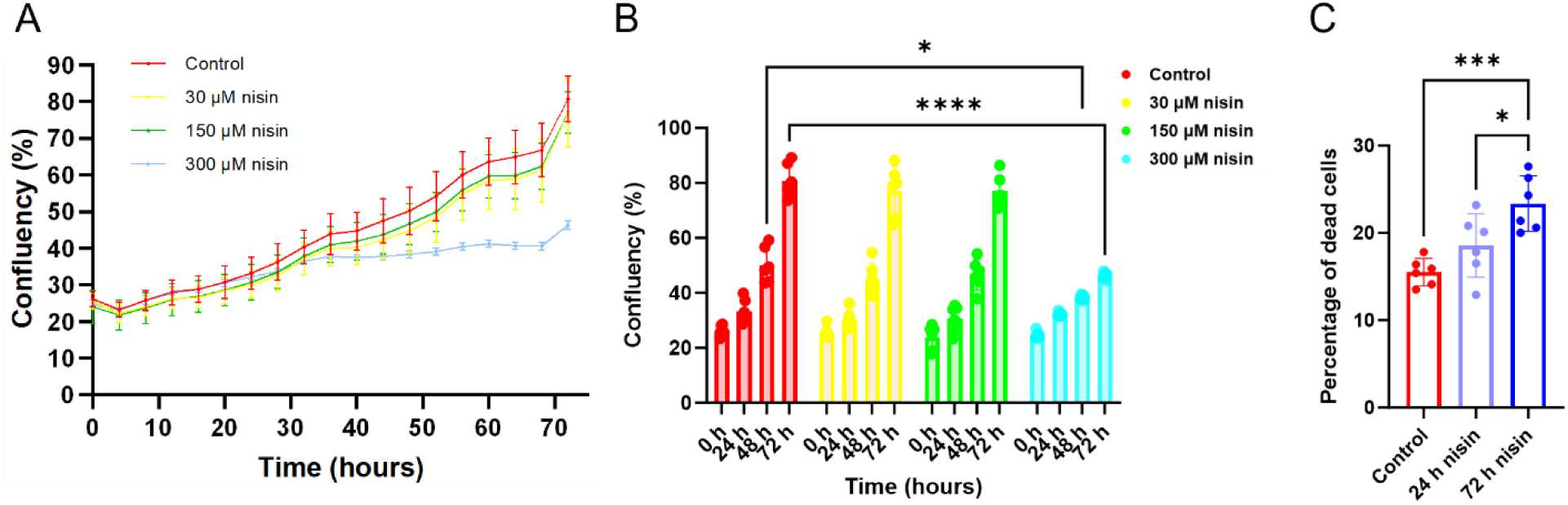
Nisin Z reduces Caco-2 cellular proliferation while increasing cell death. **(A)** The vehicle only control group, as well as cells treated with 30 µM or 150 µM nisin Z, steadily proliferate and reached ∼80% confluency at 72 h. Cells treated with 300 µM (1 mg/ml) nisin Z show reduced proliferation after 48 h of treatment. **(B)** Nisin Z significantly decreases proliferation 48 h and 72 h post treatment at 300 µM. The control group (as well as the 30 µM and 150 µM nisin Z treatments) had a ∼3.2-fold increase in confluency at 72 h, whereas the increase for 300 µM nisin Z was only ∼1.8-fold. *P* values shown were calculated using two-way ANOVA (*n* = 6 repeats; \**p* < 0.05, \*\*\*\**p* < 0.0001). **(C)** After 72 h, ∼15% of cells in the vehicle-only control group are dead. Exposure to nisin Z for 72 h slightly increases the proportion of dead cells to 23%. Statistical analysis was performed with one-way ANOVA (*n* = 6; \*\*\**p* < 0.001). Individual points represent individual experiments, bars represent group means, and error bars represent SD.

### Oral nisin Z reduces the number and size of small intestinal adenomas in Apc^Min/+^ mice

We next tested the effects of nisin Z on GI adenoma formation in the C57BL/6J Apc^Min/+^ animal model. We selected this model because Apc^Min/+^ mice are genetically predisposed to develop adenomas across their GI tract. This adenoma development can be modulated by therapeutic or environmental interventions, including modifications in the diet and microbiota^14,15^. To model an inflammatory environment that induces adenoma formation in our Apc^Min/+^ model, similar to dietary-driven inflammation-induced intestinal adenoma formation in humans, the animals were fed a high-fat diet post-weening^16,17,18^. As Apc^Min/+^ tumor formation is characterized by sex-dependent differences, we used all female animals in this study^19,20^. Nisin Z was provided to the experimental group in their drinking water for 12 weeks post weening at a concentration of 1 mg/ml (300 µM), which is within the typical range for oral antibiotics administered in animal models and corresponds to the concentration for which we observed bioactivity in our Caco-2 cell-based assay (Fig. 1). The control mice received regular drinking water. After 12 weeks on treatment, we dissected out the small and large intestines and prepared them for gross morphological analysis using our 3D-printable MIST device^21^. The nisin Z group had significantly fewer tumors in the middle region of their GI tract compared to the control animals (Fig. 2A). In addition, nisin Z treatment also resulted in a ∼50% reduction in tumor area in this region (Fig. 2B). We did not observe significant differences in tumor number or area in other small intestinal regions or the colon (Fig. 2A, B).

**Fig. 2.**
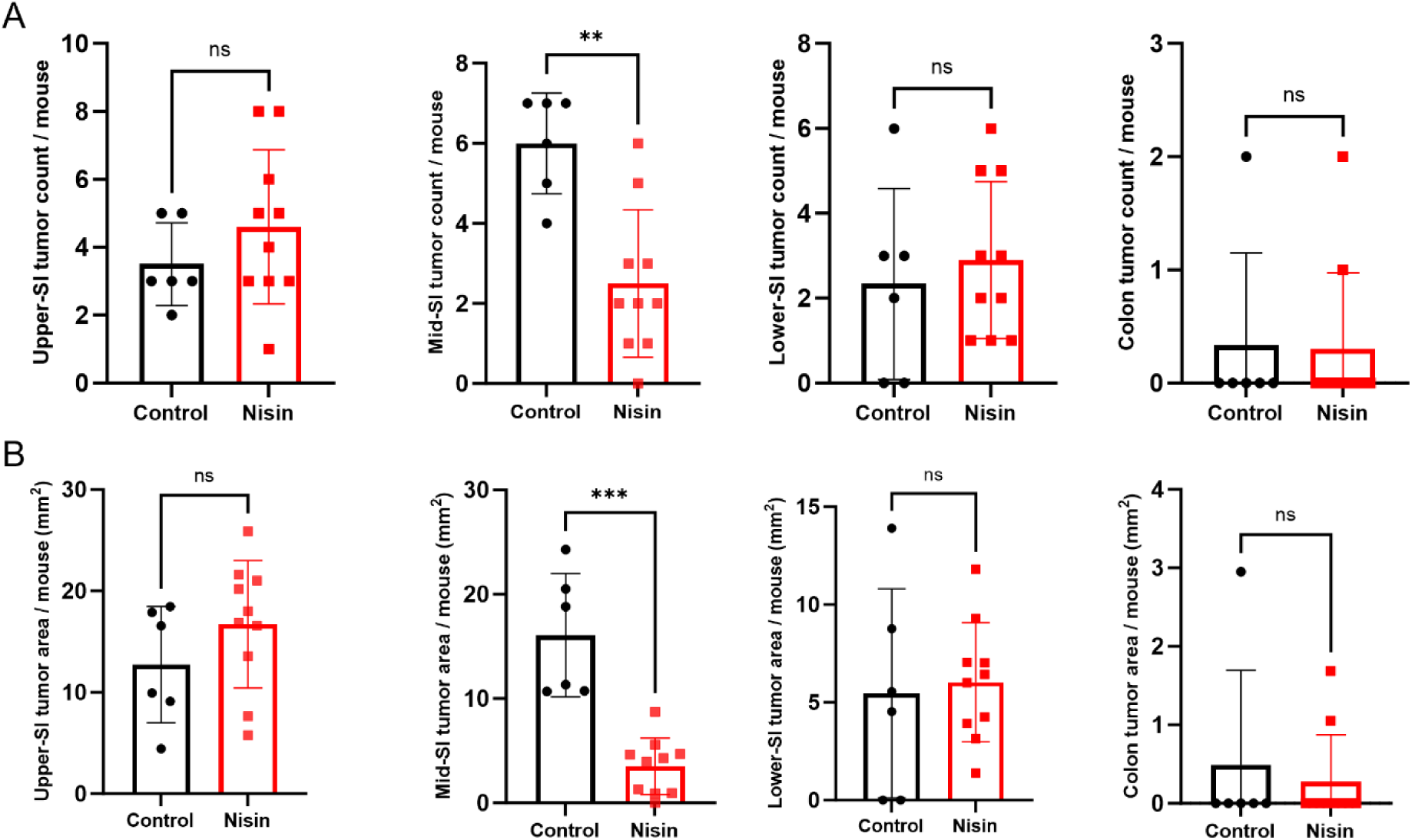
Oral administration of nisin Z in the drinking water of Apc^Min/+^ mice reduces tumor numbers and sizes in the middle region of the small intestine after 12 wks. **(A)** Tumor counts of the upper, middle, and lower small intestine (SI), as well as the colon of Apc^Min/+^ mice on regular drinking water or drinking water (control) containing 1 mg/ml nisin Z. **(B)** Tumor burden is represented by the sum of the tumor areas per mouse for each intestinal segment. Statistical analysis was performed using a Mann-Whitney test (*n* = 6-10 animals; \*\**p* < 0.01, and \*\*\**p* < 0.001). Individual points represent individual animals, bars represent group means and error bars represent SD.

Since our cell-based assays suggested that the effect of nisin Z might be mostly due to reduced proliferation, we next compared cyclin D1 expression in the middle small intestine and the colon. Cyclin D1 is a crucial cell cycle regulatory protein that permits the transition of G1 to S phase^22^. We observed a significant decrease in cyclin D1 expression in the middle region of the small intestine of nisin Z treated Apc^Min/+^ mice (Fig. 3A). Unlike the colon, where nisin Z had no effect on cyclin D1 expression (Fig. 3B). Our results suggest that nisin Z could be associated with the observed lower tumor burden in the middle section of the small intestine by decreasing cyclin D1 expression and thereby reducing tumor cell proliferation.

**Fig. 3.**
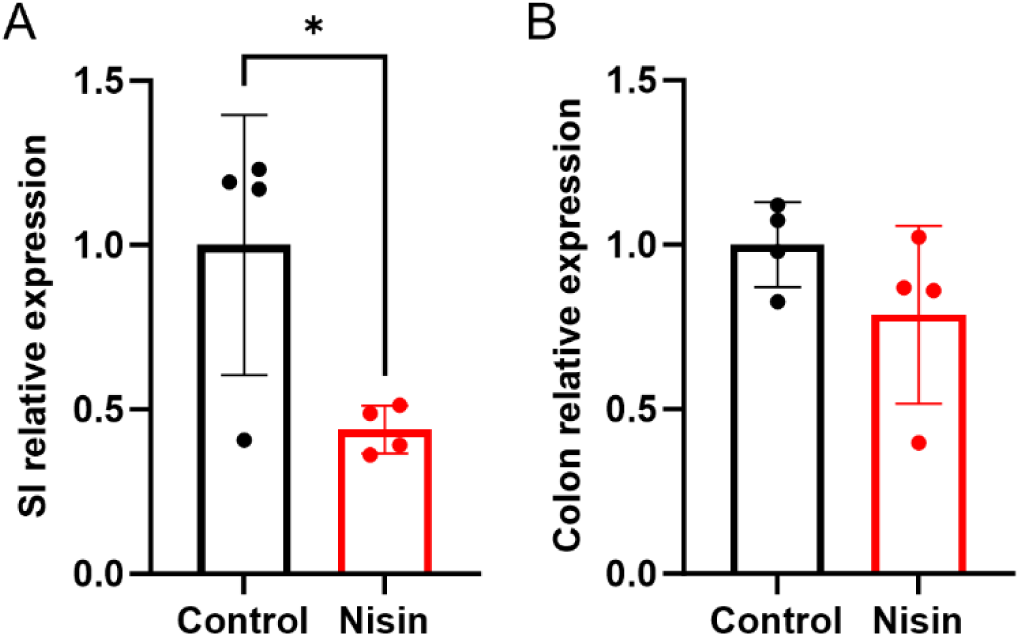
Nisin Z reduces cyclin D1 expression in the middle region of the small intestine of Apc^Min/+^ mice. **(A)** Oral nisin Z (at 1 mg/ml) reduces cyclin D1 expression in the middle region of the small intestine (SI) after 12 wks. **(B)** Cyclin D1 expression in the colon is not significantly affected by oral nisin Z treatment. Statistical analysis was performed using an unpaired t-test (*n* = 4 animals; \**p* < 0.05). mRNA expression levels were calculated based on the ΔΔ-CT method and genes of interest were normalized to the housekeeping gene β-actin. Individual points represent individual animals, bars represent group means and error bars represent SD.

### Nisin Z attenuates the NF-κB mediated inflammatory response

Since tumor development in the Apc^Min/+^ model can be affected by several factors like diet, gut microbiota, as well as an inflammatory environment, we decided to examine these different factors further. As an antibiotic, nisin Z could alter the composition of the gut microbiota, which can result in an indirect effect on the inflammatory environment in the GI tract. Alternatively, nisin Z could exert a direct, microbiota-independent anti-inflammatory effect. In Apc^Min/+^ mice, inflammation is associated with tumor progression through pathways such as the Nuclear factor kappa B (NF-κB)^23–25^. We tested the effect of nisin Z on NF-κB signaling using the human monocytic THP-1 Blue reporter assay. Addition of the Toll-like receptor 4 ligand lipopolysaccharide (LPS) results in an activation of the NF-κB pathway, which is quantified via the activity of a secreted embryonic alkaline phosphatase. Pretreatment of the cells with a range of different nisin Z concentrations (1.5 µM – 150 µM) for 4 hours prior to the LPS challenge resulted in a dose-dependent reduction in NF-κB activation (Fig. 4A). This indicates that nisin Z could have the capability of directly reducing the response to a proinflammatory stimulus, even in the absence of the gut microbiota or their metabolites. Since we observed a modest cytotoxicity effect of nisin Z on Caco-2 cells, we next investigated whether the observed decrease in NF-κB activation at higher concentrations is due to modulation of the signaling pathway rather than increased cell death. We performed a live/dead stain and found no differences in THP1 viability across the concentration range tested (Fig. 4B). Taken together, these results show that nisin Z can have a modest, direct anti-inflammatory effect, which could contribute to reducing the inflammatory environment in our animal model.

**Fig. 4.**
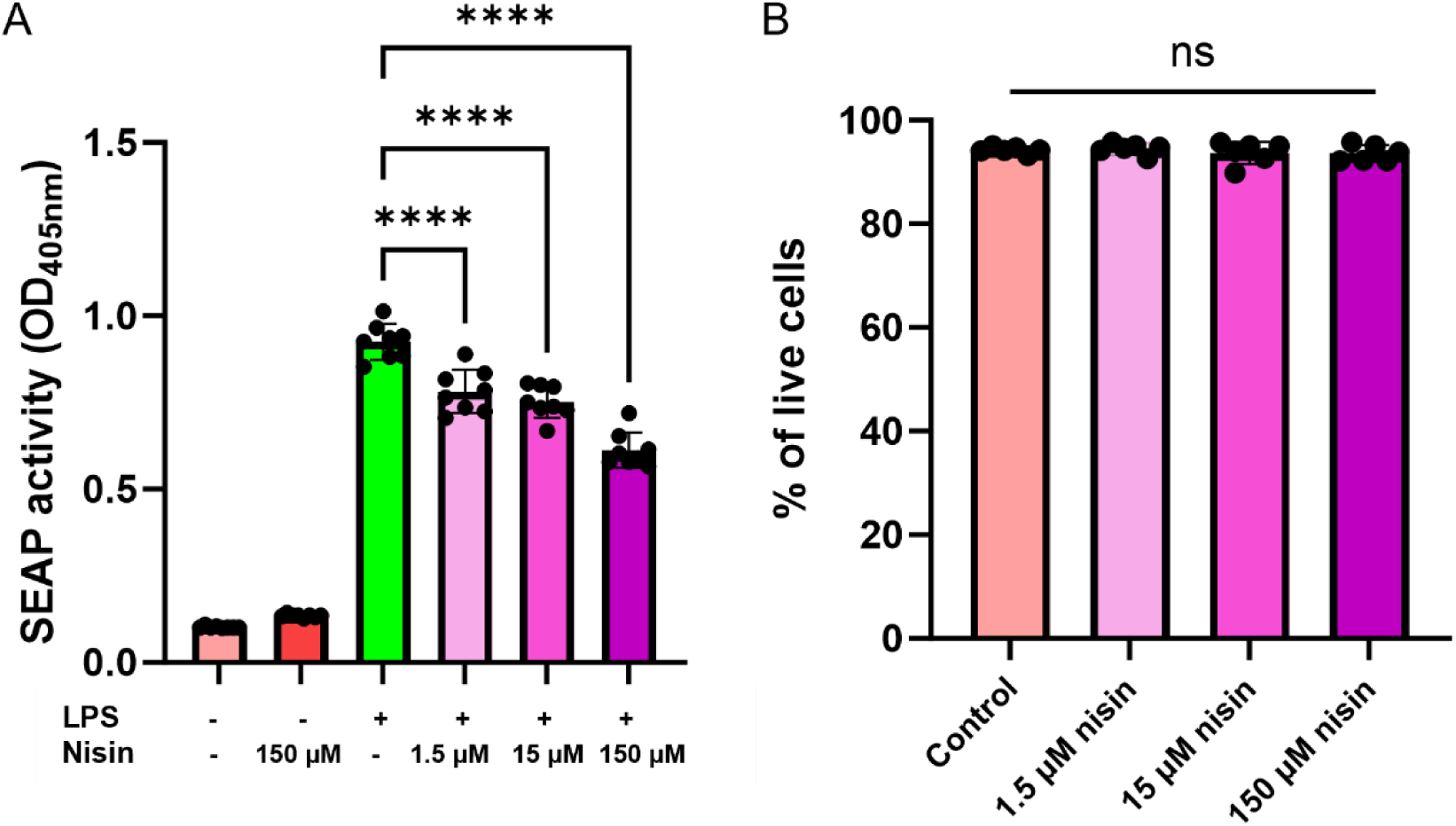
Nisin Z attenuates an LPS-induced proinflammatory response in a dose-dependent manner. **(A)** The NF-κB mediated proinflammatory response is monitored using the THP-1 Blue monocyte reporter assay. Induction of NF-κB is coupled to an increase in the secreted embryonic alkaline phosphatase (SEAP), for which the activity is quantified using a colorimetric assay. A proinflammatory response is induced with 50 ng/mL LPS, and addition of different nisin Z concentrations (1.5µM, 15µM, and 150µM). **(B)** THP-1 Blue cells maintain their viability across the range of nisin Z concentrations in the NF-κB assay. Statistical analysis was performed using one-way ANOVA with a post-hoc Dunnett’s test (*n* = 6-8 repeats; \*\*\*\**p* < 0.0001). Individual points represent individual experiments, bars represent group means and error bars represent SD.

### Oral nisin Z treatment differentially affects the cecal microbiome composition and microbiome-dependent plasma metabolites in WT and Apc^Min/+^ mice

Since nisin Z is primarily known for its antibiotic properties, we next assessed its potential indirect anti-inflammatory impact via the gut microbiota and their metabolites. We used 16S rRNA sequencing to determine the cecal microbiota composition of Apc^Min/+^ mice on nisin Z versus control drinking water. To account for potential differences due to host genotype, we compared these data to the cecal communities of C57BL/6J WT female littermates, which were randomized into similar control and nisin Z drinking water groups. To account for potential cage effects, we included cecal samples from animals that were housed in two distinct cages for each group. As can be expected from an antibiotic intervention, nisin Z reduced the alpha diversity in WT mice, as estimated by the Chao1 diversity index (Fig. 5A). Interestingly, the Apc^Min/+^ alpha-diversity was less impacted by nisin Z treatment, though a similar reduced trend was observed. Canonical correspondence analysis highlights the difference in beta-diversity between the control WT and Apc^Min/+^ communities, which seem to converge in a separate cluster upon nisin Z treatment (Fig. 5B). The total diversity plot shows general trends in response to nisin Z treatment that are observable in both the WT and Apc^Min/+^ backgrounds (Fig. 5C). Most notably, nisin Z reduces the relative abundance of *Faecalibaculum* and *Lactobacillus* at the expense of relative increases in *Helicobacter*, *Akkermansia*, *Odoribacter*, *Lachnospiraceae* (A2), *Bilophila*, and *Alistipes*. These differences are also reflected in the total diversity plot (Fig. 5D) and clustering heat map (Fig. 5E), which additionally revealed a decimation *of Romboutsia* upon nisin Z treatment, as well as underscores the contributions of *Lachnospiraceae, Faecalibaculum,* and *Lactobacillus* to community differences in Apc^Min/+^ mice. Our data show that nisin Z causes shifts in the microbial communities, as would be expected from an antibiotic, and in addition highlights potential differences in community and nisin Z response between WT and Apc^Min/+^ littermates.

**Fig. 5.**
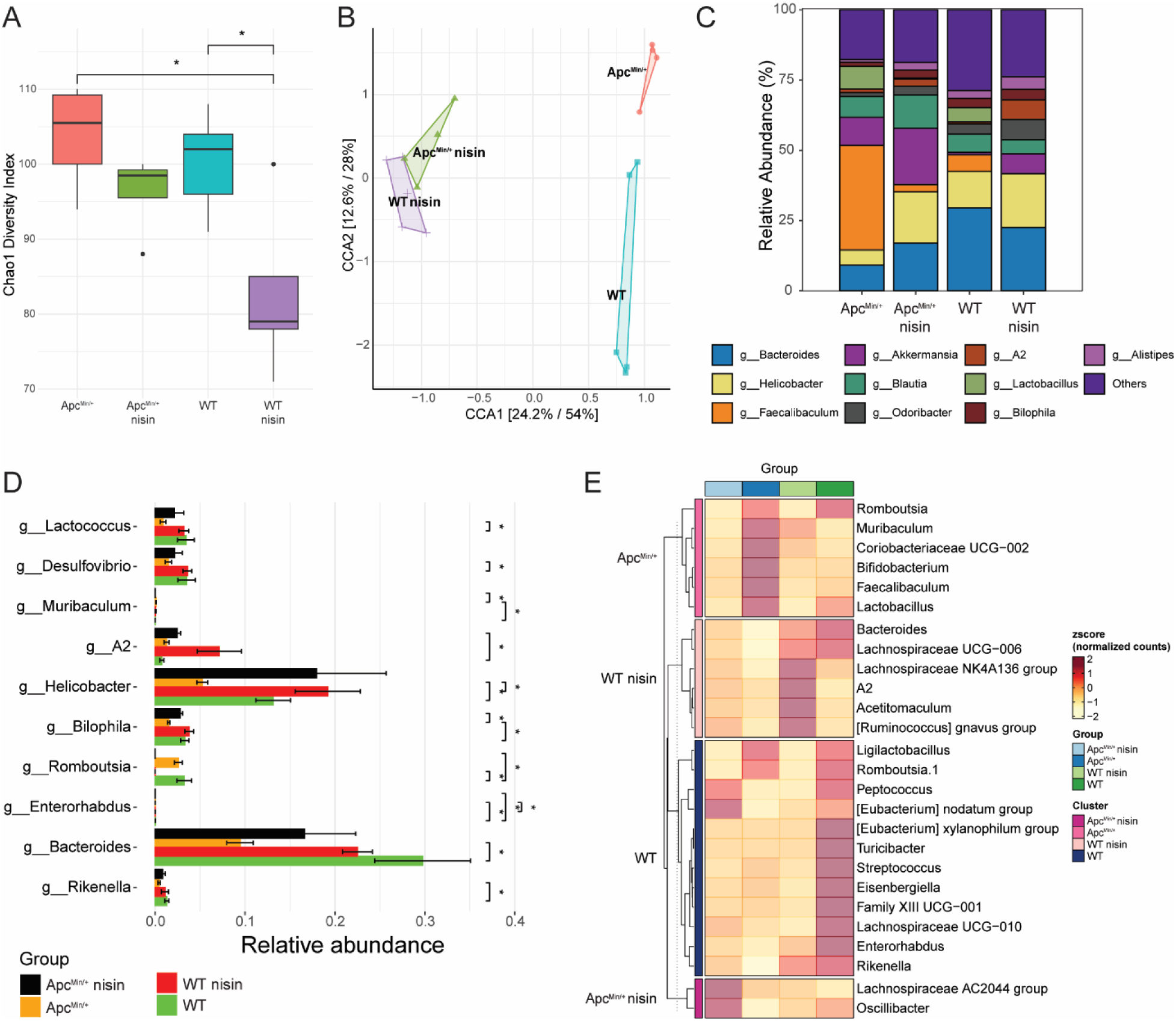
Orally administered nisin Z differentially affects the cecal microbiome composition in WT and Apc^Min/+^ mice. **(A)** Alpha-diversity (as estimated by Chao1 diversity index) is significantly more impacted by nisin Z treatment in WT animals. Pairwise analysis was done using Wilcoxon rank sum t-test. **(B)** Canonical correspondence analysis shows convergence of the beta-diversity of WT and Apc^Min/+^ mice treated with nisin, in contrast to the rather distinct control microbial communities for these mouse genotypes. We used permanova analysis (R2=0.36, p-value =0.014). **(C)** Stacked bar chart of relative abundances (left y-axis) of the top genera for each of the four groups. **(D)** Differential abundance analysis highlighting the genera that are significantly altered by nisin Z treatment or mouse genotype. Pairwise test used was “metagenomeSeq” with Benjamini Hochberg correction. **(E)** Heatmap of the differentially abundant taxa among the different groups. Statistical test used was ANOVA with Benjamini Hochberg correction The color gradient represents the relative abundance z-score (*n* = 4-5 animals; \**p* < 0.05).

To inquire whether the observed differences in gut microbiome composition could have anti-inflammatory implications, we analyzed the peripheral plasma from the WT and Apc^Min/+^ mice using a targeted LC-MS/MS panel for microbiome-associated metabolites. In line with the microbiome composition data, we observed host genotype-dependent effects on the systemic concentration of certain metabolites in response to nisin Z treatment (Fig. 6). In Apc^Min/+^ mice, the plasma concentration of the microbiome-influenced metabolite serotonin was increased with nisin Z treatment, whereas the amounts of phenyl sulfate and 4-hydroxyhippuric acid were lower compared to the control animals. In WT animals, nisin Z was found to impact the plasma concentration of hippuric acid. Interestingly, nisin Z treatment resulted in genotypic differences in the plasma concentrations of 4-hydroxyphenylpyruvic acid, indoxyl sulfate, and phenylpyruvic acid.

**Fig. 6.**
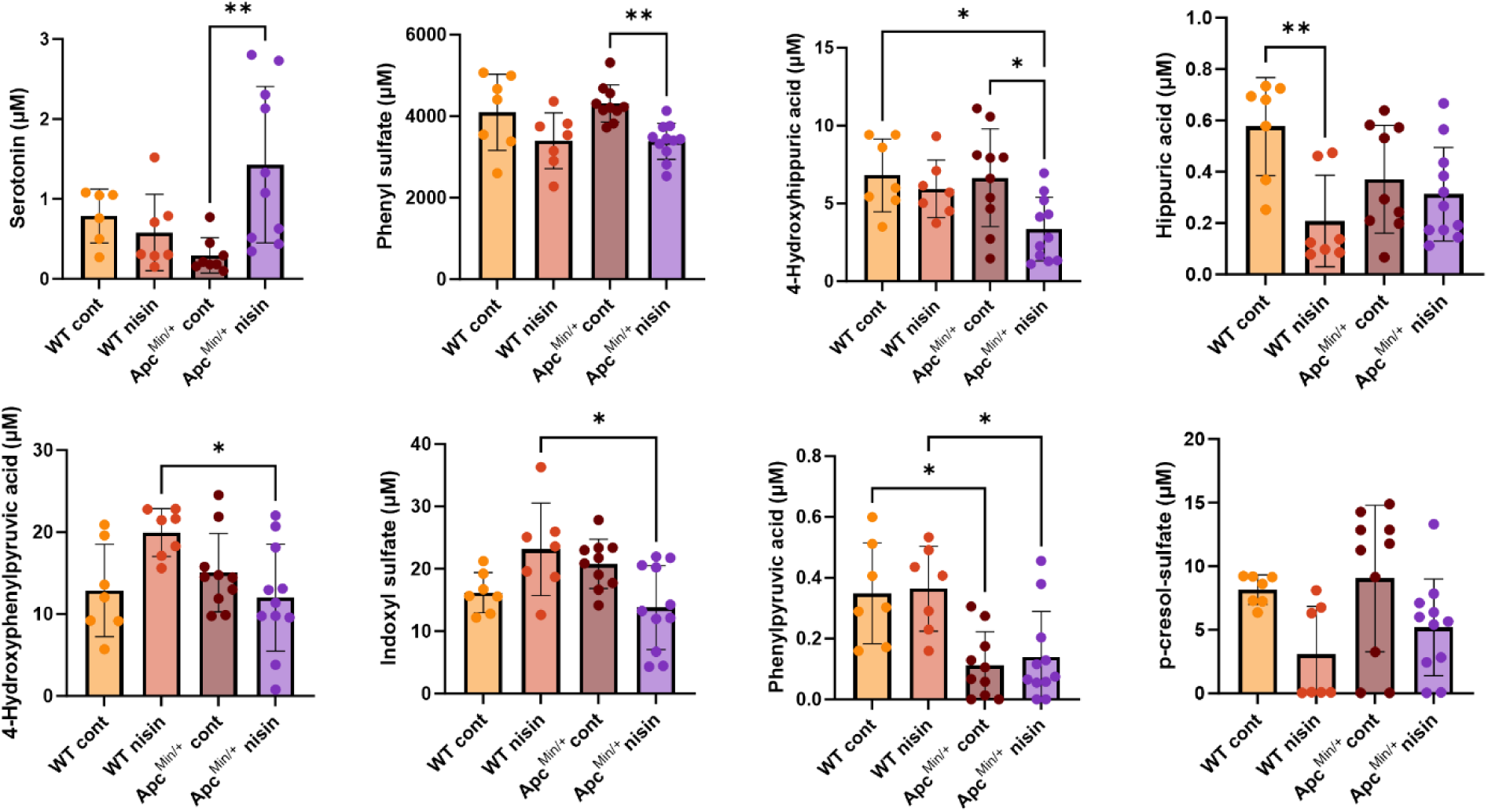
Oral nisin Z treatment differentially affects gut microbiome-dependent plasma metabolites in WT and Apc^Min/+^ mice. Peripheral plasma concentrations of microbiome-dependent metabolites were measured by LC-MS/MS, including phenyl sulfate, hippuric acid, serotonin, indoxyl sulfate, and 4-hydroxyhippuric acid. Statistical analysis was performed using the Kruskal-Wallis nonparametric test (*n* = 7-10 animals; \**p* < 0.05, \*\**p* < 0.01, and \*\*\**p* < 0.001. Individual points represent individual animals, bars represent group means and error bars represent SD.

## Discussion

Nisin Z is a post-translationally modified ribosomal peptide antibiotic produced by certain strains of *L. lactis* subsp. *lactis* and is commonly used in food preservation^26^. More recently, the compound has gained interest for its potential anti-tumor activity in head and neck cancer^9^. Additional *in vitro* studies have revealed similar anti-proliferative effects on CRC cell lines^10,11,27,28^, though no *in vivo* efficacy has yet been demonstrated for this cancer type. Our *in vitro* observations that nisin Z reduces Caco-2 proliferation and has a modest cytotoxic effect are in line with these prior studies. We also explored a potential direct anti-inflammatory effect of nisin Z via modulation of the NF-κB pathway, and this might be linked to the observed reduction in cellular proliferation^29^. NF-κB signaling has been linked to G1-S phase transition as well as inflammatory pathways, and we found that nisin Z can exhibit modest anti-inflammatory activity through this pathway. Overall, a reduction in inflammation is associated with reduced tumor formation in the Apc^Min/+^ model ^29,30^ as well as with a better overall colorectal cancer prognosis^25^.

Here we report the first *in vivo* investigation of the potential anti-intestinal adenoma activities of nisin Z using the Apc^Min/+^ animal model. We observed a decrease in the number and size of tumors in the middle section of the small intestine, which was associated with a decrease in cyclin D1 expression, indicating that a sufficient concentration of bioactive nisin Z might be present in this GI region. We did not observe a nisin Z-dependent effect on the tumor burden or cyclin D1 expression in the colon. We speculate that nisin Z might be partially degraded along the GI tract due to its proteinaceous nature, thereby not retaining sufficient concentrations to exert anti-tumor activity in the lower small intestine and colon. The apparent lack of effect in the upper small intestinal segment could potentially be due to differences in transit time and pH. Despite the potential partial degradation of nisin Z in the upper GI tract, it still had a significant impact on the cecal microbiome composition, as well as microbiome-dependent plasma metabolites. It remains to be determined whether these effects are due to a modulation of the small intestinal microbiota with downstream impact on the cecum, or whether sufficient amounts of nisin Z make it to the lower GI tract to exert antibiotic activity. Nisin was found to exert similar effects on the microbiota in a porcine model, suggesting survival through the GI tract^31^. While an earlier study of the GI transit for the related compound lacticin 3147 did not detect any intact compound across the GI tract by mass spectrometry, potentially due to a lower administered oral dose^32^.

We observed genotype-specific changes in the gut microbiota of WT and Apc^Min/+^ mice upon treatment with nisin Z. As expected from its antimicrobial activity spectrum, nisin Z causes an overall decrease in the relative abundance of Gram-positive bacteria. Several of the differentially abundant genera in our data have previously been implicated in colorectal cancer. Nisin Z treatment increased the abundance of *Bacteroides* in our Apc^Min/+^ mice. This is beneficial, as a decrease in *Bacteroides* abundance has been associated with worse CRC outcomes, though this is highly strain-dependent^33^. Additionally, we observed an increase in *Akkermansia* with nisin Z treatment, which has previously been linked to a reduction in intestinal inflammation through the NF-κB pathway, leading to a decrease in tumor formation^34^. An increase in *Odoribacter* might protect the host against CRC through enhanced cellular apoptosis^35,36^. We also observed some unexpected changes in the microbiota, which could have a negative impact on the GI inflammatory environment. In contrast to previously published work, nisin did not inhibit the growth of *Helicobacter* in our animals^37^. Certain *Helicobacter* species (such as *H. pylori*) have been associated with the development of gastric and CRC^38^, yet this is unlikely to be the case in our animals, which were raised in specific-pathogen-free conditions. Furthermore, nisin Z treatment increased the abundance of *Bilophila* in Apc^Min/+^ mice. *Bilophila* spp. are prominent producers of hydrogen sulfide, which can enhance inflammation and cellular proliferation, potentially worsening CRC^39^. Overall, the gut microbiome analysis by us, as well as others, suggest that nisin Z has anti-inflammatory properties and impacts microbial composition and diversity^31,40^.

We performed a functional interrogation of the microbiota by measuring microbiome-associated metabolites in the peripheral plasma. Apc^Min/+^ mice receiving nisin Z treatment showed increased serotonin levels. The majority of serotonin is produced in the gut from tryptophan, and this process is influenced by the gut microbiota^41^. *Lactobacillus* spp. are among these serotonin regulators, and while their abundance was previously found to be directly correlated to serotonin levels^42^, *Lactobacillus* were actually reduced in our nisin Z treated animals. We also observed a decreasing trend in the plasma concentrations of the uremic toxins, indoxyl sulfate and p-cresol sulfate, in nisin Z treated Apc^Min/+^ mice. *Bacteroides spp.* are prominent producers of indoxyl sulfate via the metabolism of tryptophan to indoles^43^. Despite a relative increase in *Bacteroides spp.* abundance, we observed a decrease in indoxyl sulfate plasma concentration, seemingly contradicting prior observations. In addition, *Coriobacteriaceae spp.* and *Clostridium spp.* have been reported to utilize tyrosine to produce p-cresol sulfate^44^. Prior studies have revealed an association between both uremic toxins and an increase in pro-inflammatory cytokines, exacerbating CRC^45–48^. This is in line with our observed direct relationship between *Coriobacteriaceae spp.* abundance and p-cresol sulfate concentration. Taken together, the reduced levels of uremic toxins could contribute to an overall decrease in luminal inflammation in nisin Z treated Apc^Min/+^ mice.

## Conclusion

We showed that nisin Z can have an anti-proliferative effect in an Apc^Min/+^ intestinal tumor mouse model via the downregulation of cyclin D expression. We speculate that the localized effect in the small intestine could be due to degradation further along the GI tract as well as differences in physiological parameters earlier in the GI tract. We did not uncover a unifying mechanism which can explain our observations, but identified direct anti-inflammatory, as well as indirect effects on the microbiota and their metabolites. We hypothesize that these individual factors, in combination with the continuous administration of nisin Z, contribute to a net reduction in the inflammatory environment, which in turn leads to the reduced small intestinal tumor burden.

## Materials and methods

### Cell culture

THP-1 Blue NF-κB Cells (InvivoGen) were cultured in customized medium and sub-cultured following the manufacturers protocol. The custom medium consists of Roswell Park Memorial Institute Medium (RPMI) 1640, 20 mM HEPES solution (Gibco), 1% GlutaMAX (Gibco), 1% Penicillin/Streptomycin, 100 µg/ml Normocin (InvivoGen), and supplemented with 10% Fetal Bovine Serum (FBS). The selection medium additionally contains 10 µg/ml blasticidin (InvivoGen). THP-1 cells were sub-cultured in selection medium every 4 days and harvested by centrifugation at 400 *x g* for 4 minutes. Supernatant was removed and cells were resuspended in fresh selection medium. Cell culture was typically maintained between 7x10^5^ – 1x10^6^ cells/ml. CRC (Caco-2) cells (ATCC, USA) were cultured according to ATCC’s recommendations. Cells were sub-cultured weekly in complete medium, consisting of Eagle’s Minimum Essential Medium (EMEM) + 20% FBS and fed 1-2 times per week. Subculturing and assays were performed at 80% confluency (1x10^5^ cells/cm^2^).

### Nisin Z solution preparation

“Ultrapure Nisin Z” (Handary S.A., Belgium) solutions were prepared fresh on the day of the experiments by vortexing for 30 seconds and sonicating for 15 minutes to ensure homogeneity. Next, solutions were passed through a 0.22 µm filter (MilliporeSigma, USA). For Caco-2 experiments, nisin Z was dissolved in complete media and for THP1-Blue experiments in custom media without blasticidin, and for animal studies, nisin Z was dissolved in drinking water.

### Caco-2 proliferation assay

Caco-2 proliferation over time was measured using an Incucyte Live-Cell Analysis System (Sartorius, USA). Caco-2 cells were seeded (4x10^4^ cells/well) and incubated overnight at 37 °C + 5% CO2, then treated with 30 µM, 150 µM, or 300 µM of nisin Z, and compared to a vehicle only control. Confluency (%) was measured at 37°C + 5% CO_2_ every 4 hours for 72 hours, using standard cell-by-cell analysis, where the software’s algorithm identifies unlabeled, discrete cells in phase images.

### NF-κB reporter assay

THP1-Blue cells (InvivoGen, USA) were seeded (1x10^4^ cells/well) into a 96-well plate and incubated overnight at 37°C + 5% CO_2_. Cells were pretreated with customized RPMI vehicle, or various concentrations of nisin Z (1.5 µM, 15 µM, 150 µM) in RPMI for 4 hours. After incubation, 50 ng/mL lipopolysaccharide (LPS, Invitrogen, USA) was added, and cells were further incubated overnight. Alkaline phosphatase produced by the cells was quantified using a *p*-Nitrophenyl Phosphate (PNPP) substrate kit (ThermoFisher, USA), according to the user guidelines. Absorbance was measured after 30 minutes at 405nm in a Biotek Synergy HT Microplate Reader (Agilent, USA).

### Cell culture live/dead assay

Caco-2 or THP-1 Blue cells were seeded at 1x10^6^ cells/well in a 6-well plate. Caco-2 cells were treated with 300 µM nisin Z for 24 h or 72 h. Next, Caco-2 cells were dissociated from the wells using TrypLE Express (Gibco, USA). THP-1 Blue cells were treated with 1.5 µM, 15 µM, or 150 µM nisin Z. Cell staining with LIVE/DEAD Fixable Blue Dead Cell Stain Kit for UV excitation (Invitrogen, USA) was performed according to the user guide. Finally, the fixed cell suspension was analyzed by flow cytometry using a FACSymphony A5 SE (BD Biosciences, USA), and the results were analyzed using FlowJo_v10.10.0 (BD Biosciences, USA).

### Animal studies

C57BL/6J WT and C57BL/6J Apc^min/+^ breeders were purchased from The Jackson Laboratory, USA, and housed in the Biological Resources Unit of the Cleveland Clinic Lerner Research Institute. At 8 weeks, C57BL/6 WT females and C57BL/6 Apc^min/+^ males were cohoused for breeding. The genotype of the pups was determined using allele-specific PCR. For experimental groups, 3-4 mice were cohoused per cage, in at least two cages per experimental group. Experimental female WT and Apc^min/+^ mice were placed on a high-fat diet (HFD) with 60 kcal% lard-derived fat (Diet# D12492, Research Diets, Inc.) at 5 weeks old. Animals were randomly assigned to groups provided with nisin Z (300 µM) or regular drinking water. Animal body weight and food intake were monitored weekly and after 12 weeks on treatment, animals were sacrificed using a ketamine (120mg/kg)—xylazine (16mg/kg) cocktail injected intraperitoneally. Peripheral plasma was collected and prepared for LC-MS/MS analysis to detect gut microbiome-related metabolites. The GI tract of the Apc^min/+^ mice was dissected, and the small intestine was sectioned into three equally sized regions: upper, middle, and lower small intestine. The small intestinal segments and colon were longitudinally sliced and spread onto a wax surface using our 3D-printed Mouse Intestinal Slicing Tool^21^. Intestinal adenomas were enumerated by two separate individuals, and their areas were measured using ImageJ (NIH, USA). Sections of the small intestine and colon were used for gene expression analysis, and cecal contents were extracted for 16S rRNA sequencing of microbiome composition. All animal studies and procedures were previously approved by the Institutional Animal Care and Use Committee at the Lerner Research Institute of the Cleveland Clinic.

### GI tissue gene expression analysis

Small intestine and colon tissue sections (50-100 mg) were lysed for 5 minutes at 30 Hz using Tissuelyser II (Qiagen, Germany). RNA was isolated using TRIzol (Invitrogen, USA) and treated with the DNase Removal Kit (Invitrogen, USA) to remove DNA contaminants. The concentration of RNA was determined by nanodrop and normalized to 750 ng/µL. cDNA was synthesized using the qScript Supermix (QuantaBio, USA) according to the manufacturer’s instructions and diluted for use with the SYBR Green Master Mix (Applied Biosystems, USA) and quantified by RT-PCR (StepOnePlus RT-PCR system, Applied Biosystems, USA). mRNA expression levels were calculated based on the ΔΔ-CT method and genes of interest were normalized to the housekeeping gene β-actin. Primers utilized: Cyclin D1, 5’-AGACCTTTGTGGCCCTCTG-3’ – F and 5’-GGCAGTCCGGGTCACACT-3’ – R. β-actin, 5’-TTCCTTCTTGGGTATGGA ATCCT-3’ – F and 5’-TTTACGGATGTCAACGTCACAC-3’.

### 16S rRNA sequencing

DNA from cecal contents was extracted using DNeasy PowerSoil Pro Kit (Qiagen, Germany). Purified DNA isolate is sequenced utilizing MiSeq Reagent Micro Kit v2 (Illumina, USA). 16S rRNA gene amplicon sequencing and bioinformatics analysis were performed using methods explained earlier (Barot et al., 2022; Chambers et al., 2022; Weiss et al., 2021). Briefly, raw 16S amplicon sequence and metadata, were demultiplexed using split_libraries_fastq.py script implemented in QIIME2 (Bolyen et al., 2019). Demultiplexed fastq file was split into sample-specific fastq files using split_sequence_file_on_sample_ids.py script from QIIME2. Individual fastq files without non-biological nucleotides were processed using Divisive Amplicon Denoising Algorithm (DADA) pipeline (Callahan et al., 2016). The output of the dada2 pipeline (feature table of amplicon sequence variants (an ASV table)) was processed for alpha and beta diversity analysis using phyloseq (McMurdie and Holmes, 2013), and microbiomeSeq (http://www.github.com/umerijaz/microbiomeSeq) packages in R. We analyzed variance (ANOVA) among sample categories while measuring the of α-diversity measures using plot_anova_diversity function in microbiomeSeq package. Permutational multivariate analysis of variance (PERMANOVA) with 999 permutations was performed on all principal coordinates obtained during CCA with the ordination function of the microbiomeSeq package. Pairwise correlation was performed between the microbiome (genera) and metabolomics (metabolites) data was performed using the microbiomeSeq package.

### Statistical analysis

Differential abundance analysis was performed using the random-forest algorithm, implemented in the DAtest package (https://github.com/Russel88/DAtest/wiki/usage#typical-workflow). Briefly, differentially abundant methods were compared with False Discovery Rate (FDR), Area Under the (Receiver Operator) Curve (AUC), Empirical power (Power), and False Positive Rate (FPR). Based on the DAtest’s benchmarking, we selected metagenomeSeq and anova as the methods of choice to perform differential abundance analysis. We assessed the statistical significance (P < 0.05) throughout, and whenever necessary, we adjusted P-values for multiple comparisons according to the Benjamini and Hochberg method to control False Discovery Rate (Benjamini and Hochberg, 1995). Linear regression (parametric test), and Wilcoxon (Non-parametric) test were performed on genera and ASVs abundances against metadata variables using their base functions in R (version 4.1.2; R Core Team, 2021) (Team, 2021).

### Gut microbiome-related metabolite LC-MS/MS analysis

Peripheral plasma samples were prepared for analysis by adding an internal standard mix in a 4:1 v/v ratio, as detailed in a previously published method^49^. Samples were vortexed and centrifuged at 20,000 x *g* for 10 mins (at 4 °C), and the supernatant was transferred to glass sample vials. LC-MS/MS analysis was performed on a LCMS-8050 Triple Quad (Shimadzu Scientific Instruments, USA) using an XSelect C18 column (2.5µM, 2.1 mm X 150mm) (Waters, Ireland). Column temperature was kept at 40 °C. Solvents A and B were 0.1% acetic acid in water and 0.1% acetic acid in acetonitrile, respectively. Samples were run according to the following gradient: 0.0-2.0min (0% B), 2.0-5.0min (40% B), 5.0-7.5min (85% B), 7.5-8.0 (100% B), 8.0-10.0min (100% B), 10.0-13.0min (0% B), and with a flow rate of 0.4 mL/min. Data was analyzed using Lab Solution software (Shimadzu Scientific Instruments, USA).

## Data availability statement

All data are presented in the manuscript or available on request. Raw sequence files from the 16S rRNA gene sequencing were deposited in Zenodo’s Sequence Read Archive (DOI 10.6084/m9.figshare.28990487.v1).

## Acknowledgements

This work was supported by seed funding from the Cleveland Clinic Foundation (J.C.), and in part by a Research Grant from the Prevent Cancer Foundation (PCF2019-JC), and a National Institutes of Health (NIH) grant R01 AI153173 (J.C.). I.N. was in part supported by the NIH R01HL16074 grant.

## Author contributions

L.H.N. and J.C. conceived and designed the experiments. L.H.N., S.A., R.L.M., B.D., V.B., and J.C. performed the mouse experiments. L.H.N., and J.E. performed the biochemical workup and transcriptional analysis of cell and mouse experiments. N.S. conducted the microbial sequencing analysis. L.H.N., S.A., and I.N. performed LC-MS/MS analysis of *in vivo* samples. L.H.N., and J.C. designed experiments and or analyzed the subsequent data. L.H.N., and J.C. wrote the manuscript and handled visualization. All authors discussed the results and commented on the manuscript.

## References

(1) Colorectal Cancer Statistics | How Common Is Colorectal Cancer? https://www.cancer.org/cancer/types/colon-rectal-cancer/about/key-statistics.html

(2) What Is Colorectal Cancer? | How Does Colorectal Cancer Start? https://www.cancer.org/cancer/types/colon-rectal-cancer/about/what-is-colorectal-cancer.html

(3) Valle, L.; Vilar, E.; Tavtigian, S. V.; Stoffel, E. M. Genetic Predisposition to Colorectal Cancer: Syndromes, Genes, Classification of Genetic Variants and Implications for Precision Medicine. J Pathol 2019, 247 (5), 574–588. 10.1002/path.5229.

(4) Murphy, N.; Moreno, V.; Hughes, D. J.; Vodicka, L.; Vodicka, P.; Aglago, E. K.; Gunter, M. J.; Jenab, M. Lifestyle and Dietary Environmental Factors in Colorectal Cancer Susceptibility. Molecular Aspects of Medicine 2019, 69, 2–9. 10.1016/j.mam.2019.06.005.

(5) Fodde, R. The APC Gene in Colorectal Cancer. Eur J Cancer 2002, 38 (7), 867–871. 10.1016/s0959-8049(02)00040-0.

(6) Ren, J.; Sui, H.; Fang, F.; Li, Q.; Li, B. The Application of ApcMin/+ Mouse Model in Colorectal Tumor Researches. J Cancer Res Clin Oncol 2019, 145 (5), 1111–1122. 10.1007/s00432-019-02883-6.

(7) Shin, J. M.; Gwak, J. W.; Kamarajan, P.; Fenno, J. C.; Rickard, A. H.; Kapila, Y. L. Biomedical Applications of Nisin. J Appl Microbiol 2016, 120 (6), 1449–1465. 10.1111/jam.13033.

(8) Khazaei Monfared, Y.; Mahmoudian, M.; Caldera, F.; Pedrazzo, A. R.; Zakeri-Milani, P.; Matencio, A.; Trotta, F. Nisin Delivery by Nanosponges Increases Its Anticancer Activity against *in-Vivo* Melanoma Model. Journal of Drug Delivery Science and Technology 2023, 79, 104065. 10.1016/j.jddst.2022.104065.

(9) Kamarajan, P.; Hayami, T.; Matte, B.; Liu, Y.; Danciu, T.; Ramamoorthy, A.; Worden, F.; Kapila, S.; Kapila, Y. Nisin ZP, a Bacteriocin and Food Preservative, Inhibits Head and Neck Cancer Tumorigenesis and Prolongs Survival. PLoS One 2015, 10 (7), e0131008. 10.1371/journal.pone.0131008.

(10) Norouzi, Z.; Salimi, A.; Halabian, R.; Fahimi, H. Nisin, a Potent Bacteriocin and Anti-Bacterial Peptide, Attenuates Expression of Metastatic Genes in Colorectal Cancer Cell Lines. Microb Pathog 2018, 123, 183–189. 10.1016/j.micpath.2018.07.006.

(11) Ahmadi, S.; Ghollasi, M.; Hosseini, H. M. The Apoptotic Impact of Nisin as a Potent Bacteriocin on the Colon Cancer Cells. Microbial Pathogenesis 2017, 111, 193–197. 10.1016/j.micpath.2017.08.037.

(12) Joo, N. E.; Ritchie, K.; Kamarajan, P.; Miao, D.; Kapila, Y. L. Nisin, an Apoptogenic Bacteriocin and Food Preservative, Attenuates HNSCC Tumorigenesis via CHAC 1. Cancer Med 2012, 1 (3), 295–305. 10.1002/cam4.35.

(13) University of California, San Francisco. A Phase I&#x2F;IIa, Single-Arm, Dose-Confirmation and Dose-Expansion Study Evaluating Changes in the Oral Microbiome of Patients With Oral Cavity Squamous Cell Carcinoma (OSCC) After Short-Term Ingestion of Nisin, a Naturally Occurring Food Preservative; Clinical trial registration NCT06097468; clinicaltrials.gov, 2025. https://clinicaltrials.gov/study/NCT06097468 (accessed 2025-05-02).

(14) Baltgalvis, K. A.; Berger, F. G.; Peña, M. M. O.; Davis, J. M.; Carson, J. A. The Interaction of a High-Fat Diet and Regular Moderate Intensity Exercise on Intestinal Polyp Development in ApcMin/+ Mice. Cancer Prev Res (Phila) 2009, 2 (7), 641–649. 10.1158/1940-6207.CAPR-09-0017.

(15) Li, L.; Li, X.; Zhong, W.; Yang, M.; Xu, M.; Sun, Y.; Ma, J.; Liu, T.; Song, X.; Dong, W.; Liu, X.; Chen, Y.; Liu, Y.; Abla, Z.; Liu, W.; Wang, B.; Jiang, K.; Cao, H. Gut Microbiota from Colorectal Cancer Patients Enhances the Progression of Intestinal Adenoma in Apcmin/+ Mice. EBioMedicine 2019, 48, 301–315. 10.1016/j.ebiom.2019.09.021.

(16) Park, M.-Y.; Kim, M. Y.; Seo, Y. R.; Kim, J.-S.; Sung, M.-K. High-Fat Diet Accelerates Intestinal Tumorigenesis Through Disrupting Intestinal Cell Membrane Integrity. J Cancer Prev 2016, 21 (2), 95–103. 10.15430/JCP.2016.21.2.95.

(17) Yang, J.; Wei, H.; Zhou, Y.; Szeto, C.-H.; Li, C.; Lin, Y.; Coker, O. O.; Lau, H. C. H.; Chan, A. W. H.; Sung, J. J. Y.; Yu, J. High-Fat Diet Promotes Colorectal Tumorigenesis Through Modulating Gut Microbiota and Metabolites. Gastroenterology 2022, 162 (1), 135–149.e2. 10.1053/j.gastro.2021.08.041.

(18) Doerner, S. K.; Reis, E. S.; Leung, E. S.; Ko, J. S.; Heaney, J. D.; Berger, N. A.; Lambris, J. D.; Nadeau, J. H. High-Fat Diet-Induced Complement Activation Mediates Intestinal Inflammation and Neoplasia, Independent of Obesity. Mol Cancer Res 2016, 14 (10), 953–965. 10.1158/1541-7786.MCR-16-0153.

(19) Cherukuri, D. P.; Ishikawa, T.; Chun, P.; Catapang, A.; Elashoff, D.; Grogan, T. R.; Bugni, J.; Herschman, H. R. Targeted Cox2 Gene Deletion in Intestinal Epithelial Cells Decreases Tumorigenesis in Female, but Not Male, Apc/+ Mice. Molecular Oncology 2014, 8 (2), 169–177. 10.1016/j.molonc.2013.10.009.

(20) McAlpine, C. A.; Barak, Y.; Matise, I.; Cormier, R. T. Intestinal-Specific PPARγ Deficiency Enhances Tumorigenesis in ApcMin/+ Mice. International Journal of Cancer 2006, 119 (10), 2339–2346. 10.1002/ijc.22115.

(21) DeLucia, B.; Samorezov, S.; Zangara, M. T.; Markley, R. L.; Osborn, L. J.; Schultz, K. B.; McDonald, C.; Claesen, J. A 3D-printable Device Allowing Fast and Reproducible Longitudinal Preparation of Mouse Intestines. Animal Model Exp Med 2022, 5 (2), 189–196. 10.1002/ame2.12228.

(22) Sherr, C. J. D-Type Cyclins. Trends Biochem Sci 1995, 20 (5), 187–190. 10.1016/s0968-0004(00)89005-2.

(23) Chronic epithelial NF-κB activation accelerates APC loss and intestinal tumor initiation through iNOS up-regulation | PNAS. https://www.pnas.org/doi/full/10.1073/pnas.1211509109 (accessed 2025-05-02).

(24) Yu, S.; Yin, Y.; Wang, Q.; Wang, L. Dual Gene Deficient Models of ApcMin/+ Mouse in Assessing Molecular Mechanisms of Intestinal Carcinogenesis. Biomedicine & Pharmacotherapy 2018, 108, 600–609. 10.1016/j.biopha.2018.09.056.

(25) Zhao, H.; Wu, L.; Yan, G.; Chen, Y.; Zhou, M.; Wu, Y.; Li, Y. Inflammation and Tumor Progression: Signaling Pathways and Targeted Intervention. Sig Transduct Target Ther 2021, 6 (1), 1–46. 10.1038/s41392-021-00658-5.

(26) de Vos, W. M.; Mulders, J. W.; Siezen, R. J.; Hugenholtz, J.; Kuipers, O. P. Properties of Nisin Z and Distribution of Its Gene, NisZ, in Lactococcus Lactis. Appl Environ Microbiol 1993, 59 (1), 213–218. 10.1128/aem.59.1.213-218.1993.

(27) Hosseini, S. S.; Goudarzi, H.; Ghalavand, Z.; Hajikhani, B.; Rafeieiatani, Z.; Hakemi-Vala, M. Anti-Proliferative Effects of Cell Wall, Cytoplasmic Extract of Lactococcus Lactis and Nisin through down-Regulation of Cyclin D1 on SW480 Colorectal Cancer Cell Line. Iran J Microbiol 2020, 12 (5), 424–430. 10.18502/ijm.v12i5.4603.

(28) Soans, S. H.; Chonche, M. J.; Sharan, K.; Srinivasan, A.; Archer, A. C. Apoptotic and Anti-Inflammatory Effect of Nisin-Loaded Sodium Alginate-Gum Arabic Nanoparticles against Colon Cancer Cells. International Journal of Biological Macromolecules 2025, 305, 141747. 10.1016/j.ijbiomac.2025.141747.

(29) Khan, A.; Zhang, Y.; Ma, N.; Shi, J.; Hou, Y. NF-ΚB Role on Tumor Proliferation, Migration, Invasion and Immune Escape. Cancer Gene Ther 2024, 31 (11), 1599–1610. 10.1038/s41417-024-00811-6.

(30) McClellan, J. L.; Davis, J. M.; Steiner, J. L.; Day, S. D.; Steck, S. E.; Carmichael, M. D.; Murphy, E. A. Intestinal Inflammatory Cytokine Response in Relation to Tumorigenesis in the Apc(Min/+) Mouse. Cytokine 2012, 57 (1), 113–119. 10.1016/j.cyto.2011.09.027.

(31) O’Reilly, C.; Grimaud, G. M.; Coakley, M.; O’Connor, P. M.; Mathur, H.; Peterson, V. L.; O’Donovan, C. M.; Lawlor, P. G.; Cotter, P. D.; Stanton, C.; Rea, M. C.; Hill, C.; Ross, R. P. Modulation of the Gut Microbiome with Nisin. Sci Rep 2023, 13 (1), 7899. 10.1038/s41598-023-34586-x.

(32) Gardiner, G. E.; Rea, M. C.; O’Riordan, B.; O’Connor, P.; Morgan, S. M.; Lawlor, P. G.; Lynch, P. B.; Cronin, M.; Ross, R. P.; Hill, C. Fate of the Two-Component Lantibiotic Lacticin 3147 in the Gastrointestinal Tract. Appl Environ Microbiol 2007, 73 (21), 7103–7109. 10.1128/AEM.01117-07.

(33) Xie, X.; He, Y.; Li, H.; Yu, D.; Na, L.; Sun, T.; Zhang, D.; Shi, X.; Xia, Y.; Jiang, T.; Rong, S.; Yang, S.; Ma, X.; Xu, G. Effects of Prebiotics on Immunologic Indicators and Intestinal Microbiota Structure in Perioperative Colorectal Cancer Patients. Nutrition 2019, 61, 132–142. 10.1016/j.nut.2018.10.038.

(34) Ma, X.; LvjunYan; Yu, X.; Guo, H.; He, Y.; Wen, S.; Yu, T.; Wang, W. The Alleviating Effect of *Akkermansia Muciniphila* PROBIO on AOM/DSS-Induced Colorectal Cancer in Mice and Its Regulatory Effect on Gut Microbiota. Journal of Functional Foods 2024, 114, 106091. 10.1016/j.jff.2024.106091.

(35) Oh, B. S.; Choi, W. J.; Kim, J.-S.; Ryu, S. W.; Yu, S. Y.; Lee, J.-S.; Park, S.-H.; Kang, S. W.; Lee, J.; Jung, W. Y.; Kim, Y.-M.; Jeong, J.-H.; Lee, J. H. Cell-Free Supernatant of Odoribacter Splanchnicus Isolated From Human Feces Exhibits Anti-Colorectal Cancer Activity. Front Microbiol 2021, 12, 736343. 10.3389/fmicb.2021.736343.

(36) Xing, C.; Du, Y.; Duan, T.; Nim, K.; Chu, J.; Wang, H. Y.; Wang, R.-F. Interaction between Microbiota and Immunity and Its Implication in Colorectal Cancer. Front Immunol 2022, 13, 963819. 10.3389/fimmu.2022.963819.

(37) Charest, A. M.; Reed, E.; Bozorgzadeh, S.; Hernandez, L.; Getsey, N. V.; Smith, L.; Galperina, A.; Beauregard, H. E.; Charest, H. A.; Mitchell, M.; Riley, M. A. Nisin Inhibition of Gram-Negative Bacteria. Microorganisms 2024, 12 (6), 1230. 10.3390/microorganisms12061230.

(38) Ralser, A.; Dietl, A.; Jarosch, S.; Engelsberger, V.; Wanisch, A.; Janssen, K. P.; Middelhoff, M.; Vieth, M.; Quante, M.; Haller, D.; Busch, D. H.; Deng, L.; Mejías-Luque, R.; Gerhard, M. Helicobacter Pylori Promotes Colorectal Carcinogenesis by Deregulating Intestinal Immunity and Inducing a Mucus-Degrading Microbiota Signature. 2023. 10.1136/gutjnl-2022-328075.

(39) Waqas, M.; Halim, S. A.; Ullah, A.; Ali, A. A. M.; Khalid, A.; Abdalla, A. N.; Khan, A.; Al-Harrasi, A. Multi-Fold Computational Analysis to Discover Novel Putative Inhibitors of Isethionate Sulfite-Lyase (Isla) from Bilophila Wadsworthia: Combating Colorectal Cancer and Inflammatory Bowel Diseases. Cancers (Basel) 2023, 15 (3), 901. 10.3390/cancers15030901.

(40) Huang, F.; Teng, K.; Liu, Y.; Wang, T.; Xia, T.; Yun, F.; Zhong, J. Nisin Z Attenuates Lipopolysaccharide-Induced Mastitis by Inhibiting the ERK1/2 and P38 Mitogen-Activated Protein Kinase Signaling Pathways. Journal of Dairy Science 2022, 105 (4), 3530–3543. 10.3168/jds.2021-21356.

(41) Strandwitz, P. Neurotransmitter Modulation by the Gut Microbiota. Brain Res 2018, 1693 (Pt B), 128–133. 10.1016/j.brainres.2018.03.015.

(42) Hizay, A.; Dag, K.; Oz, N.; Comak-Gocer, E. M.; Ozbey-Unlu, O.; Ucak, M.; Keles-Celik, N. *Lactobacillus Acidophilus* Regulates Abnormal Serotonin Availability in Experimental Ulcerative Colitis. Anaerobe 2023, 80, 102710. 10.1016/j.anaerobe.2023.102710.

(43) Devlin, A. S.; Marcobal, A.; Dodd, D.; Nayfach, S.; Plummer, N.; Meyer, T.; Pollard, K. S.; Sonnenburg, J. L.; Fischbach, M. A. Modulation of a Circulating Uremic Solute via Rational Genetic Manipulation of the Gut Microbiota. Cell Host Microbe 2016, 20 (6), 709–715. 10.1016/j.chom.2016.10.021.

(44) Saito, Y.; Sato, T.; Nomoto, K.; Tsuji, H. Identification of Phenol- and p-Cresol-Producing Intestinal Bacteria by Using Media Supplemented with Tyrosine and Its Metabolites. FEMS Microbiol Ecol 2018, 94 (9), fiy125. 10.1093/femsec/fiy125.

(45) Ribeiro, A.; Liu, F.; Srebrzynski, M.; Rother, S.; Adamowicz, K.; Wadowska, M.; Steiger, S.; Anders, H.-J.; Schmaderer, C.; Koziel, J.; Lech, M. Uremic Toxin Indoxyl Sulfate Promotes Macrophage-Associated Low-Grade Inflammation and Epithelial Cell Senescence. Int J Mol Sci 2023, 24 (9), 8031. 10.3390/ijms24098031.

(46) Ichisaka, Y.; Yano, S.; Nishimura, K.; Niwa, T.; Shimizu, H. Indoxyl Sulfate Contributes to Colorectal Cancer Cell Proliferation and Increased EGFR Expression by Activating AhR and Akt. Biomed Res 2024, 45 (2), 57–66. 10.2220/biomedres.45.57.

(47) Dalal, N.; Jalandra, R.; Bayal, N.; Yadav, A. K.; Harshulika; Sharma, M.; Makharia, G. K.; Kumar, P.; Singh, R.; Solanki, P. R.; Kumar, A. Gut Microbiota-Derived Metabolites in CRC Progression and Causation. J Cancer Res Clin Oncol 2021, 147 (11), 3141–3155. 10.1007/s00432-021-03729-w.

(48) Al Hinai, E. A.; Kullamethee, Piyarach; Rowland, Ian R.; Swann, Jonathan; Walton, Gemma E.; and Commane, D. M. Modelling the Role of Microbial P-Cresol in Colorectal Genotoxicity. Gut Microbes 2019, 10 (3), 398–411. 10.1080/19490976.2018.1534514.

(49) Nemet, I.; Li, X. S.; Haghikia, A.; Li, L.; Wilcox, J.; Romano, K. A.; Buffa, J. A.; Witkowski, M.; Demuth, I.; König, M.; Steinhagen-Thiessen, E.; Bäckhed, F.; Fischbach, M. A.; Tang, W. H. W.; Landmesser, U.; Hazen, S. L. Atlas of Gut Microbe-Derived Products from Aromatic Amino Acids and Risk of Cardiovascular Morbidity and Mortality. Eur Heart J 2023, 44 (32), 3085–3096. 10.1093/eurheartj/ehad333.

